# Rapid presynaptic maturation in naturally regenerating axons

**DOI:** 10.1101/2022.06.09.493421

**Authors:** Lorcan P. Browne, Andres Crespo, Matthew S. Grubb

## Abstract

Successful neuronal regeneration requires the re-establishment of synaptic connectivity. Crucial to this process is the reconstitution of presynaptic machinery responsible for controlling neurotransmitter release. In the mammalian adult CNS post-injury regeneration is usually only possible after extensive experimental intervention, and it is unknown how presynaptic function is re-established, let alone how it might be optimised to promote functional recovery. Here we addressed these questions by studying presynaptic maturation during a regenerative process that occurs entirely naturally. After toxin-induced injury, olfactory sensory neurons in the adult mouse olfactory epithelium can regenerate fully, sending axons to the brain to re-establish synaptic contact with postsynaptic partners in the olfactory bulb. Using electrophysiological recordings in acute slices, we found that after initial re-contact, functional connectivity in this system was rapidly established. Moreover, re-connecting presynaptic terminals had almost mature functional properties, including high release probability and a strong capacity for presynaptic inhibition. Release probability then matured quickly, rendering re-established terminals functionally indistinguishable from controls just one week after initial contact. These data show that successful synaptic regeneration in the adult mammalian brain is not quite a ‘plug-and-play’ process; instead, almost-mature presynaptic terminals undergo a rapid phase of functional maturation to re-integrate into established target networks.

## Introduction

For successful recovery to occur after damage to the nervous system, re-grown or replaced axons must establish functional connections with appropriate downstream synaptic partners. This requires, amongst other critical phenomena, the maturation of presynaptic specialisations for neurotransmitter-containing vesicle release. In the peripheral nervous system of amphibians and mammals, where spontaneous axonal regeneration and behavioural recovery can occur after nerve injury, re-connecting motor neuron terminals initially display immature presynaptic properties. These include lower quantal content, lower release probability, altered calcium dynamics and fragmented active zones, which combine to produce small, sub-threshold end-plate potentials in response to nerve stimulation (Argentieri et al., 1992; Carmignoto et al., 1983; Decino, 1981; Dennis and Miledi, 1974; Ko, 1984). Subsequent maturation however, permits complete functional recovery of postsynaptic responses (Dennis and Miledi, 1974). This contrasts with presynaptic mechanisms in successfully regenerating lamprey spinal cord, where behavioural recovery can be established despite fully mature axon terminals possessing abnormal release characteristics (Oliphint et al., 2010; Parker, 2017). And in C. elegans, DA9 axon re-growth and the successful re-formation of axon terminals in appropriate locations are not accompanied by the maturation of functional release machinery, with resultingly poor behavioural recovery (Ding and Hammarlund, 2018). Even when anatomical axonal recovery is spontaneous and successful, then, it can still be accompanied by a range of different presynaptic outcomes, and there is no guarantee that functional connections will be re-established.

In the injured adult mammalian CNS, where axon re-growth or replacement is only possible with considerable experimental intervention, nothing is currently known about how presynaptic terminals re-connect with downstream circuits. Here even the most successful anatomical regeneration is usually accompanied by incomplete functional synaptic or behavioural recovery, suggestive of a disconnect between experimentally-induced axon regrowth and re-connectivity (Anderson et al., 2018; Ceto et al., 2020; Cheah et al., 2016; De Virgiliis et al., 2020; Fawcett, 2020; Lim et al., 2016; Lu et al., 2012; Wang et al., 2015). Indeed, it was recently shown that certain components of presynaptic release machinery act as a brake on axon regrowth after mammalian spinal cord injury, meaning that molecular mechanisms for promoting growth and those responsible for subsequent functional integration might be diametrically opposed (Hilton et al., 2021). But when even persuading re-growing mammalian central axons to reach their targets is already extremely difficult, how can we determine the additional manipulations required to promote their subsequent re-connectivity?

One approach is to learn lessons from a unique axonal projection in the adult mammalian CNS that is capable of regeneration. Olfactory sensory neurons (OSNs) in the nasal olfactory epithelium project axons to the brain, where they terminate in the glomerular layer of the olfactory bulb (OB). After even widespread peripheral damage due to infection, inflammation, toxicity or injury, this entire nose-to-brain axonal projection is capable of natural regeneration throughout life, with OSN axons regrowing, re-connecting with postsynaptic OB neurons, and supporting the recovery of olfactory-driven behaviour (Blanco-Hernández et al., 2012; Cheung et al., 2013; Graziadei and Graziadei, 1979; Huang et al., 2021; Schwob et al., 2017). The re-establishment of the glomerular olfactory map during this process has been well described – overall, this can be very precise but is less accurate when the initial damage to the projection is more severe (Blanco-Hernández et al., 2012; Cheung et al., 2013; Christensen et al., 2001; Costanzo, 2000; Cummings et al., 2000; Gogos et al., 2000; St John et al., 2003). We also know that, under non-injured conditions of ongoing constitutive replacement, immature OSN axon terminals can make highly dynamic, activity-dependent functional synaptic contacts with OB neurons (Cheetham et al., 2016; Huang et al., 2021). However, the functional presynaptic properties of re-connecting OSN axons remain entirely unstudied. Under baseline conditions, OSN presynaptic terminals have specialised features, characterised by high release probability and strong presynaptic inhibition via D_2_ and GABA_B_ receptors (McGann, 2013; Murphy et al., 2004; Vaaga et al., 2017). These features are already present during early postnatal development, or in the very first stages of OSN contact onto immature adult-born OB interneurons (Grubb et al., 2008). But are post-injury, initially re-connecting OSN terminals also already fully functionally mature?

Here we use an olfactotoxin model of OSN degeneration and subsequent natural regeneration to study the functional maturation of re-connecting presynaptic terminals in the adult mammalian CNS. We find that initially re-connecting OSN-to-OB inputs have almost fully mature functional properties, even at the very first stages of anatomical recovery. However, they do go through a brief maturational period where they have slightly lower release probability than control terminals. Our findings show that naturally regenerating axon terminals have many mature features as soon as they reach their targets, but nevertheless undergo a rapid process of functional development as they integrate into established post-synaptic circuitry.

## Results

### Anatomical regeneration of OSN-to-OB inputs

We induced degeneration and subsequent natural regeneration of olfactory sensory neurons (OSNs) using a single intraperitoneal injection of the selective olfactotoxin methimazole (MMZ) in adult wild-type mice (Fig 1A; Bergman and Brittebo, 1999; Blanco-Hernández et al., 2012; Huang et al., 2021; Kikuta et al., 2015). Immunohistochemical labelling for the mature OSN marker olfactory marker protein (OMP) revealed the efficacy and time course of this process in the sensory periphery. The nasal olfactory epithelium was almost entirely devoid of OMP-positive neurons 5 days post-MMZ injection (DPI), before regeneration and maturation of new-born OSNs (Graziadei and Graziadei, 1979; Schwob et al., 2017) produced an increase in mature OSN density starting from ~10 DPI (Fig 1B,C). This replenishment of the OSN population continued steadily for the following 11 weeks, reaching control levels by 90 DPI (Fig 1B,C).

**Figure 1.**
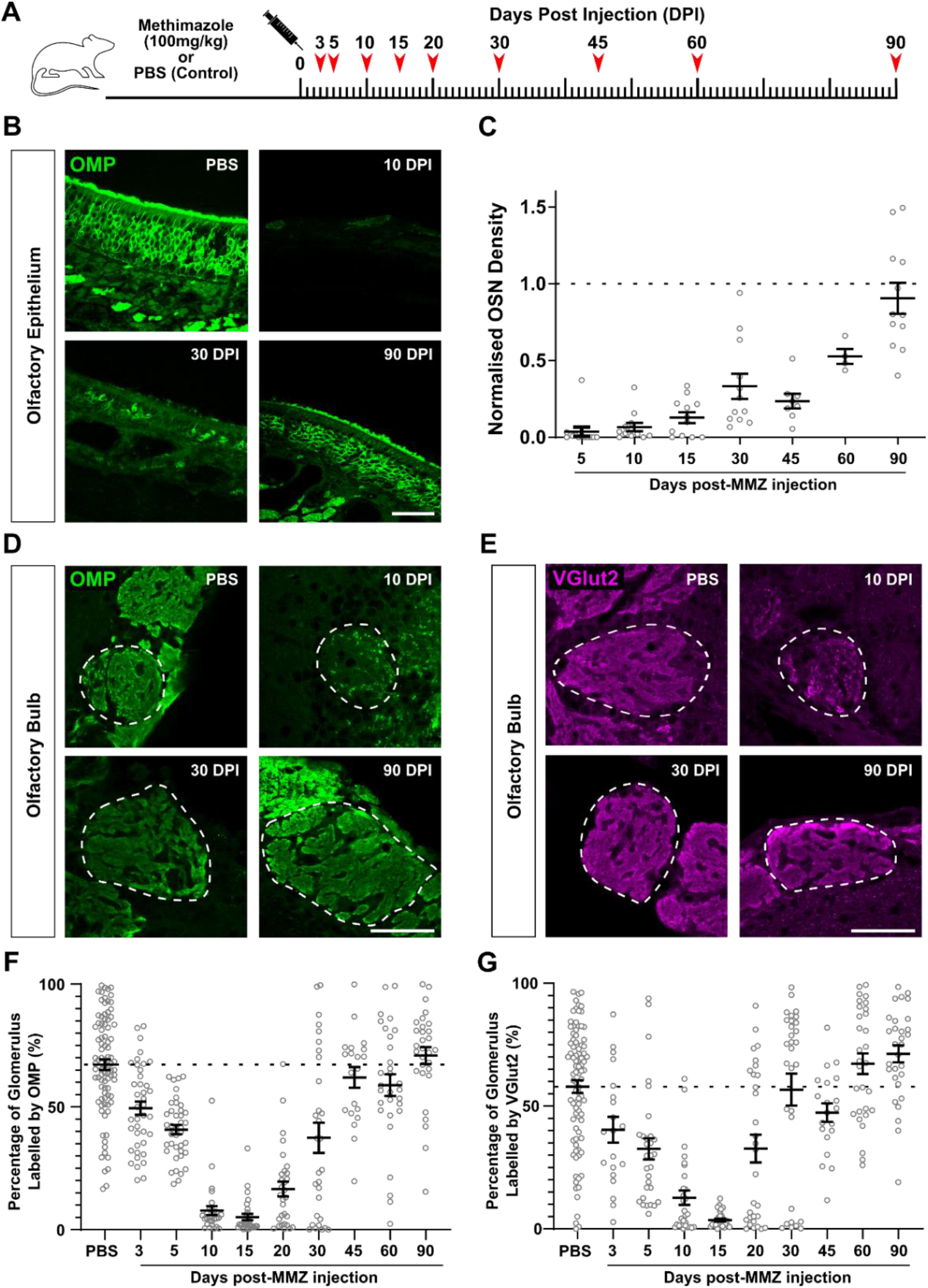
Time course of anatomical regeneration of OSN-to-OB inputs. ***A***. Schematic representation of the experimental strategy. ***B***. Example single-plane confocal images of mature OSNs labelled with OMP immunofluorescence in OE sections from Control (PBS-) or MMZ-injected mice at various timepoints post-MMZ injection. Scale bars, 50μm. ***C***. Quantification of OMP+ cell density in the OE at different timepoints post injection. All counts were normalised to values obtained from a co-embedded control slice (defined as 1; dotted line; see Methods). Dots show data from individual images; lines show mean ± SEM. ***D, E.*** Example single-plane confocal images of OMP-(***D***) or VGlut2- (***E***) stained sections of the OB taken from Control (PBS-) or MMZ-injected mice at various timepoints post-injection. Scale bars, 50μm. ***F, G***. Quantification of the glomerular coverage of OMP and VGlut2 label (see Methods) at different timepoints after MMZ injection. Dots show individual glomeruli; lines show mean ± SEM; dotted line shows control mean.

We also established the post-MMZ time course of anatomical degeneration and regeneration of OSN axon terminals in the glomerular layer of the olfactory bulb (OB). We started with immunohistochemical labelling for OMP, which is distributed throughout OSN axons (Fig 1D). Quantitative analysis of OMP-positive label coverage within the glomerular neuropil (see Methods) showed that full degeneration of distal OSN axon components in the brain took longer than the degeneration of OSN somas in the nose, only reaching minimum values at ~10 DPI (Fig 1D,F). After this, regeneration began quickly, with OMP-positive glomerular coverage increasing rapidly again from 15 DPI and regaining control values by 45 DPI (Fig 1D,F). We saw a great deal of inter-glomerular variability in this anatomical regeneration phase – at 30 DPI, for example, some glomeruli had recovered full levels of OMP innervation while others were still almost entirely denervated (Fig 1F). This variability decreased as full anatomical recovery was reached.

To specifically study the time course of OSN axon terminal degeneration and regeneration after MMZ treatment, we stained for the presynaptic vesicle-associated protein VGlut2, which is found specifically and selectively in OSN axon terminals in the OB’s glomerular layer (Fig 1E; Gabellec et al., 2007). Quantitative analysis of VGlut2 glomerular coverage revealed a very similar pattern to that of OMP, with maximal degeneration at ~15 DPI, regeneration from ~15-60 DPI, and large inter-glomerular variability at intermediate phases (Fig 1E,G). Given the reported late maturation of VGlut2 expression in OSNs (Marcucci et al., 2009), it was interesting to note that anatomical recovery of VGlut2 glomerular coverage did not lag behind, and if anything slightly preceded that of OMP during MMZ-induced regeneration (Fig 1G).

### Functional regeneration of OSN-to-OB inputs

In order to relate this time course of anatomical degeneration and regeneration to the functional properties of OSN presynaptic terminals, we prepared acute OB slices from MMZ-treated and control mice for electrophysiological recordings (Fig 2A,B). The presynaptic function of OSN terminals is well maintained in this preparation (McGann et al., 2005; Murphy et al., 2004), and can be studied by pairing electrical stimulation of OSN axons in the olfactory nerve layer with whole-cell patch-clamp recordings from OB postsynaptic neurons. We chose to record from a specialised type of excitatory interneuron in the glomerular layer – external tufted cells (ETCs) – which receive strong, reliable monosynaptic glutamatergic input from OSN axons (Hayar et al., 2004), and which can be readily identified for recording in wild-type slices based on their size, soma shape, position, and intrinsic functional properties (Galliano et al., 2021; Hayar et al., 2004). By including a cytoplasmic dye in the patch pipette, we filled all recorded ETCs to assess their anatomical integrity after the slicing procedure (Fig 2C). Of those with intact dendritic tufts extending into the glomerular neuropil, almost all experienced sharp-onset, low-latency, presumed monosynaptic evoked post-synaptic currents (EPSCs) in response to ONL stimulation (92 %, 42/50 cells; Fig 2C,D). Such responses, however, were never seen in ETCs whose dendritic tufts had been severed by the slicing process (0/61 cells; Fig 2C,D). After MMZ treatment, we were still able to detect some anatomically intact ETCs with functional OSN inputs up to 7 DPI, in agreement with the slow anatomical degeneration of OSN axons and terminals in the OB (Fig 1). However, between ~7-11 DPI we were unable to evoke any monosynaptic EPSCs in intact ETCs, despite assaying a range of stimulus locations in the ONL and a large range of stimulus intensities. In step with the anatomical recovery of OSN terminals (Fig 1), we started to see OSN inputs to ETCs re-emerge at ~13 DPI (Fig 2D). Thereafter the rate of anatomical re-connection was surprisingly rapid, and by 18 DPI the percentage of intact ETCs receiving monosynaptic OSN inputs was already back to control levels (Fig 2D).

**Figure 2.**
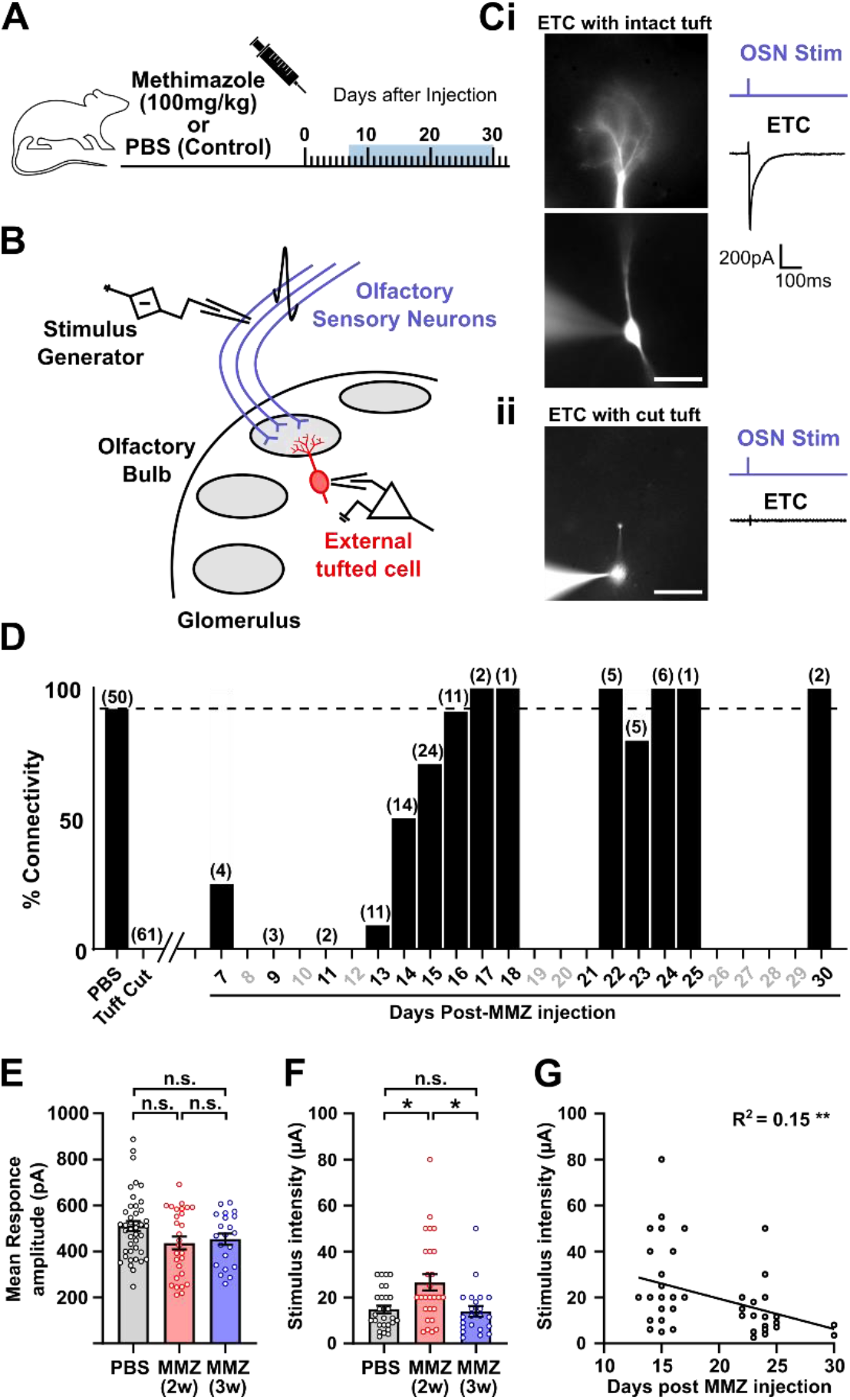
Time course of functional regeneration of OSN-to-OB inputs. ***A,B***. Schematic representation of the experimental strategy. Whole-cell patch clamp recordings were obtained from external tufted cells (ETCs) during electrical stimulation of OSN axons, in acute OB slices from mice injected with either PBS or methimazole (MMZ). Experiments were carried out at a range of timepoints from 7 DPI to 30 DPI. ***C***. Epifluorescent image showing an intact Alexa Fluor-filled ETC during a whole-cell patch-clamp recording (i). Electrical stimulation of nearby OSN axons reliably produces monosynaptic EPSCs in these cells. In cases where the ETC dendrite was severed in the slicing process, no evoked EPSC was ever observed (ii). ***D***. Plot showing the probability of successfully recording a monosynaptic OSN-to-ETC connection at different post-MMZ timepoints. Days in bold show timepoints where recordings from intact ETCs took place; numbers in brackets show the number of ETCs recorded. ***E,F***. Plots showing EPSC amplitude measured from ETCs *(**E***) and current stimulus amplitude employed to evoke the EPSC responses (***F***). 2w, 2 weeks post-MMZ; 3w, 3 weeks post-MMZ; dots show individual cells; bars show mean ± SEM; n.s., non-significant; *, P < 0.05. ***G***. Correlation plot showing the decline in current stimulus amplitude required to produce consistent response amplitudes between 2 and 3 weeks post-MMZ. Dots show individual cells; line and values show results of best-fit linear regression; **, P <0.01.

Given the varied and unpredictable routes taken by OSN axons to reach their glomerular targets (Potter et al., 2001) it is impossible to stimulate all of the inputs to a given glomerulus to reliably measure the amplitude of evoked EPSCs. However, we did find that initially re-connecting inputs from ~13-20 DPI required stronger stimulation intensities to produce reliable monosynaptic EPSCs in our target amplitude range of 200-900 pA (Fig 2E-G; stimulation current threshold; PBS, mean ± SEM, 14.8 ± 1.6 μA, n = 28 cells; MMZ(2w), 26.6 ± 3.6 μA, n = 27 cells; MMZ(3w), 14.0 ± 2.3 μA, n = 22 cells; Kruskal-Wallis test, K-W statistic = 9.66, P = 0.008; Mean rank diff, PBS vs. MMZ (2w) −14.7 μA, P = 0.043; PBS vs. MMZ (3w), +3.5 μA, P > 0.999; MMZ (2w) vs. MMZ (3w) +18.2 μA, P = 0.013, Dunn’s multiple comparisons test). Initial functional connectivity is therefore established rather quickly, with each ETC receiving at least *some* OSN inputs by a very early stage in the post-MMZ regeneration process. However, that initial input is weaker, with full functional re-innervation taking at least a week longer (Fig 1F,G).

### Newly formed OSN-to-OB synapses have a transiently higher paired-pulse ratio

Based on our characterisations of anatomical and functional OSN regeneration, we then focused on the initial period of re-connectivity between 2-3 weeks post-MMZ when OSN terminals are just beginning to re-establish contact with their post-synaptic targets in the OB (Fig 3A). Are the presynaptic terminals of re-connecting axons functionally mature at this time? We started to address this question using a paired-pulse stimulation paradigm, in which the relative amplitude of the response to the second of a pair of closely spaced stimuli can give an indication of presynaptic release probability (P_r_). Usually, a lower paired-pulse ratio (PPR), in which the second response is relatively smaller, is reflective of higher P_r_ (Dobrunz and Stevens, 1997). Indeed, at OSN-to-OB synapses in control conditions, where P_r_ is known to be unusually high, the PPR is correspondingly low (Murphy et al., 2004) and changing P_r_ by manipulating [Ca^2+^]_e_ produces reliable PPR alterations (Grubb et al., 2008). Accordingly, for inter-stimulus intervals (ISIs) of either 50 ms or 500 ms we saw low PPR values below 1 in our control slices (Fig 3B), indicative of high P_r_ under baseline conditions. In slices from 2 week post-MMZ mice, PPR values were less than 1, indicative of reasonably high P_r_ even at the earliest stage of functional re-connectivity. However, PPR at this time was significantly higher than in controls for both 50 ms and 500 ms ISIs (Fig 3B,C,D,F; 50ms ISI PPR; PBS, mean ± SEM, 0.67 ± 0.02, n = 43 cells; MMZ(2w), 0.76 ± 0.03, n = 27 cells; MMZ(3w), 0.64 ± 0.02, n = 21 cells; Kruskal-Wallis test, K-W statistic = 13.24, P = 0.0013; Mean rank diff, PBS vs. MMZ (2w) −18.8, P = 0.011, Dunn’s multiple comparisons test; 500 ms ISI PPR; PBS, mean ± SEM, 0.80 ± 0.02, n = 40 cells; MMZ(2w), 0.86 ± 0.01, n = 25 cells; MMZ(3w), 0.85 ± 0.01, n = 18 cells; Kruskal-Wallis test, K-W statistic = 6.852, P = 0.033; Mean rank diff, PBS vs. MMZ (2w) −14.8, P = 0.049, Dunn’s multiple comparisons test). By 3 weeks post-MMZ, PPR values for 50 ms ISIs had returned to control levels, and we saw a significant negative correlation of this measure with regeneration time (Fig 3D,E; Mean rank diff, PBS vs. MMZ (3w), +7.2, P = 0.92; MMZ (2w) vs. MMZ (3w) +26.0, P = 0.002, Dunn’s multiple comparisons test; Spearman correlation of 50ms ISI PPR vs MMZ DPI, Slope −0.013 ± 0.003, n = 48 cells, R^2^ = 0.24, P < 0.001). For 500 ms ISIs, PPR values remained high at 3 weeks post-MMZ, though not significantly higher than control (Fig 3F; Mean rank diff, PBS vs. MMZ (3w), −12.4, P = 0.21; MMZ (2w) vs. MMZ (3w) +2.4, P > 0.999, Dunn’s multiple comparisons test), and we saw no significant correlation between this measure and time post-MMZ injection (Fig 3G; Spearman correlation of 500 ms ISI PPR vs MMZ DPI, Slope −0.002 ± 0.002, n = 48 cells, R^2^ = 0.024, P = 0.32). These data suggest that in the earliest stages of re-connection with their postsynaptic targets regenerating OSN terminals already have rather high P_r_, which then rapidly increases to reach even higher control levels.

**Figure 3.**
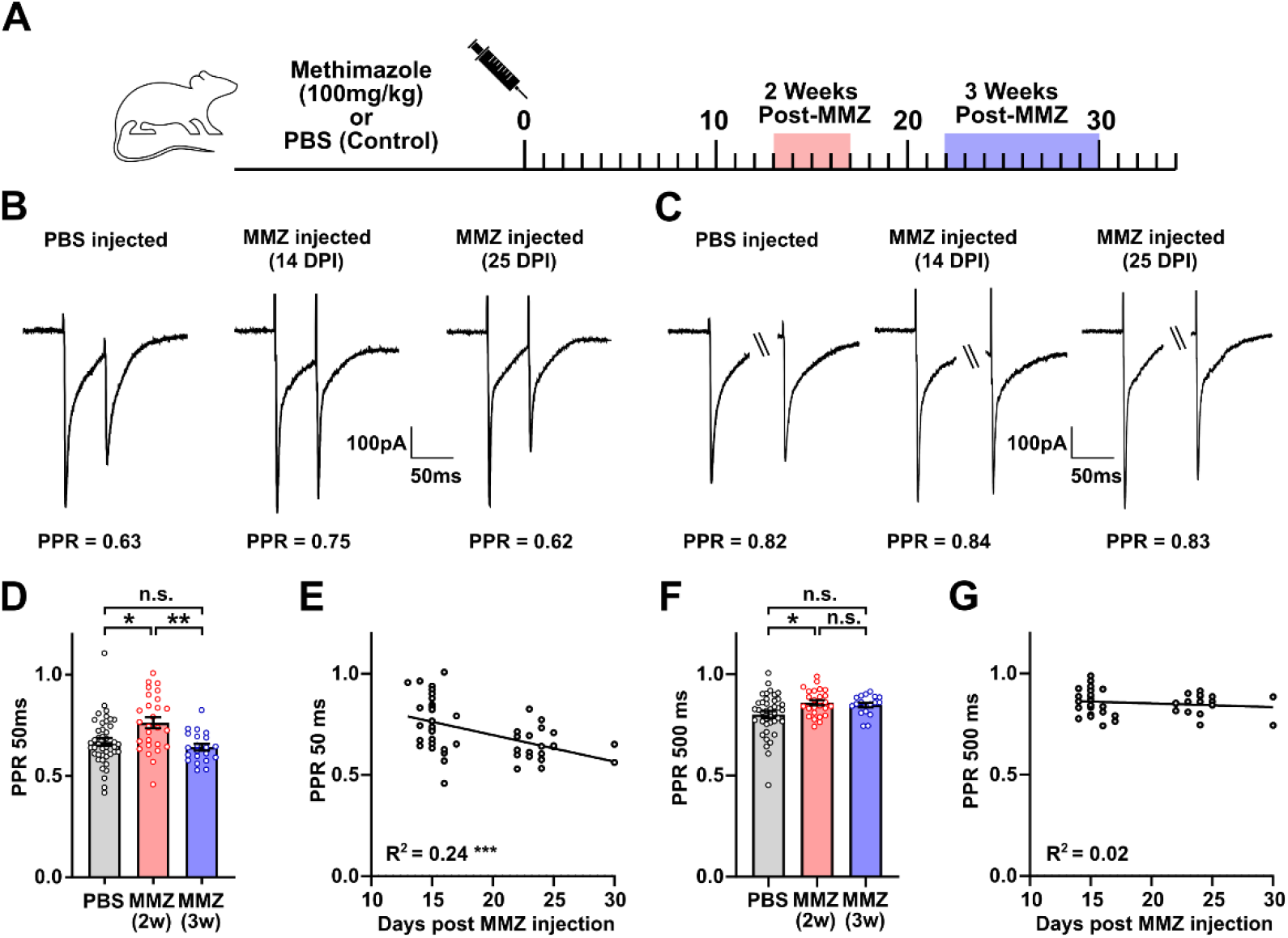
Newly formed OSN-to-OB synapses have transiently higher paired-pulse ratio. ***A***. Schematic representation of the experimental strategy to examine the properties of newly arriving (2 weeks post-methimazole (MMZ); pink shade) and maturing (3 weeks post-MMZ; blue shade) OSN axons. ***B***. Representative postsynaptic currents recorded in ETCs in response to a paired-pulse OSN stimulus with 50 ms inter-stimulus interval (ISI), in PBS- (left) or MMZ-injected mice at 14 (middle) or 25 (right) days post-injection (DPI). PPR, paired-pulse ratio. ***C***. Representative responses in the same cells to a paired-pulse stimulus with 500 ms ISI. Traces have been cropped at the break marks to show individual responses more clearly. ***D***. Plot showing the maturation of PPR with 50 ms ISI. 2w, 2 weeks post-MMZ; 3w, 3 weeks post-MMZ; dots show individual cells; bars show mean ± SEM; *, P < 0.05; **, P < 0.01; n.s., non-significant. ***E***. Correlation plot showing the decline in 50 ms ISI PPR between 2 and 3 weeks post-MMZ. Dots show individual cells; line and values show results of best-fit linear regression; ***, P < 0.001; ***F***. Maturation of PPR with 500 ms ISI. All conventions as in ***D***. ***G***. Correlation plot shows a much weaker relationship between 500 ms ISI PPR and days post-MMZ injection. All conventions as in ***E***.

### Postsynaptic receptor desensitisation has no effect on PPR

Responses to paired-pulse stimuli, however, can be influenced by many factors. Post-synaptic phenomena can also contribute to short-term synaptic depression, so we wanted to assess their potential contribution to our findings of increased PPR during OSN re-connection, starting with the passive properties of our recorded ETCs. After MMZ treatment we found a small but significant decrease in ETC whole-cell membrane capacitance and a trend towards increased input resistance, potentially suggestive of a slight decrease in total ETC membrane area during OSN regeneration (cell capacitance, PBS, mean ± SEM 51.2 ± 2.0 pF, n = 37 cells; MMZ(2w), 44.3 ± 2.2 pF, n = 26 cells; MMZ(3w), 46.3 ± 2.0 pF, n = 21 cells; one way ANOVA, F = 3.17, P = 0.047; Mean diff, PBS vs. MMZ (2w), +6.9 pF, P = 0.054; PBS vs. MMZ (3w), +4.8 pF, P = 0.35; MMZ (2w) vs. MMZ (3w), −2.1 pF, P > 0.999, Bonferroni’s multiple comparisons test; input resistance, PBS, 139.6 ± 11.4 MΩ, n = 37 cells; MMZ(2w), 176.0 ± 21.9 MΩ, n = 26 cells; MMZ(3w), 164.7 ± 18.8 MΩ, n = 21 cells; Kruskal Wallis test, K-W statistic = 2.306, P = 0.32). However, these effects are unable to explain the differences we observed in PPR, since on a cell-by-cell basis neither passive property was significantly correlated with the degree of paired-pulse depression recorded (Spearman correlation of cell capacitance vs 50 ms ISI PPR, R^2^ < 0.001, P = 0.82, n = 80; Spearman correlation of input resistance vs 50 ms ISI PPR, R^2^ = 0.002, P = 0.64, n = 80).

Postsynaptic AMPA receptor desensitisation can also contribute to PPR (Trussell et al., 1993), so we recorded OSN-evoked EPSCs before and after the application of 100 μM cyclothiazide to block this process. As previously reported for OSN-to-OB synapses (Murphy et al., 2004), blocking AMPA receptor desensitisation increased response amplitudes without altering 50 ms PPR (Fig 4). These effects were no different between control and 2 week post-MMZ slices (Fig 4; % first peak amplitude after cyclothiazide application, PBS, mean ± SEM, 136.2 ± 8.2 %, n = 13 cells; MMZ(2w), 119.0 ± 4.2 %, n = 10 cells; unpaired t-test with Welch’s correction, t = 1.87, df = 17.5, P = 0.079; % 50 ms ISI PPR after cyclothiazide application, PBS, mean ± SEM, 104.5 ± 4.1 %, n = 13 cells; MMZ(2w), 109.1 ± 3.1 %, n = 10 cells; Unpaired t test with Welch’s correction, t = 0.89, df = 20.6, P = 0.38), indicating that post-synaptic factors are unlikely to account for the increased PPR we observe in regenerating OSN inputs.

**Figure 4.**
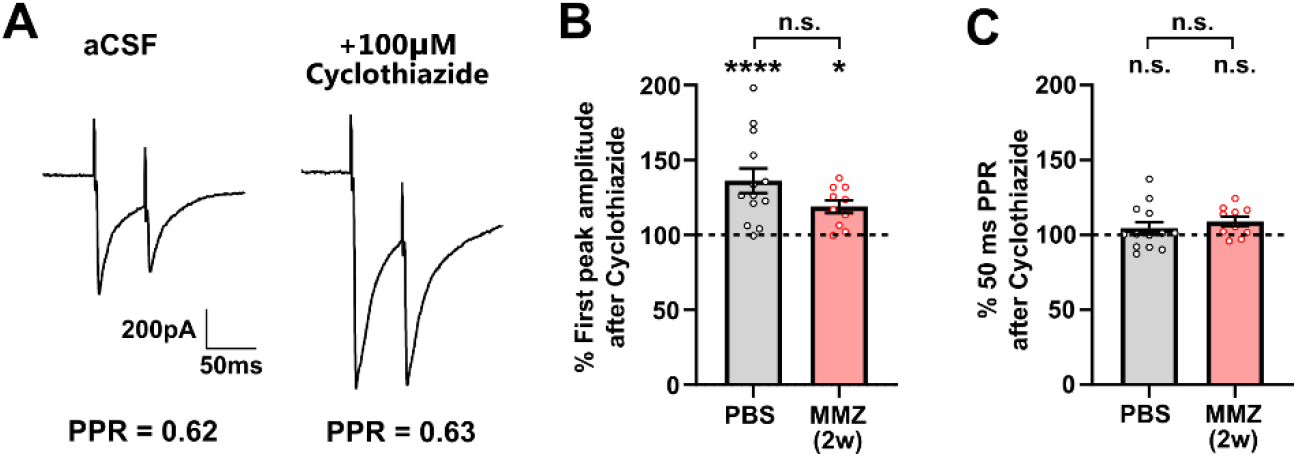
Postsynaptic receptor desensitisation has no effect on PPR at re-connecting OSN-to-OB synapses. ***A**.* Representative EPSCs measured in an ETC in response to a paired-pulse OSN stimulus with 50 ms inter-stimulus interval, before (left, aCSF) and after (right) application of 100μM cyclothiazide to block AMPA receptor desensitisation. PPR, paired-pulse ratio. ***B***. Mean ± SEM % change in OSN-to-ETC first peak amplitude in PBS- and MMZ-injected (2w, 2 weeks post-injection) animals after application of 100μM cyclothiazide. Dots show individual cells; *, P < 0.05; ****, P < 0.0001; n.s., non-significant. ***C***. Mean ±SEM % change in OSN-to-ETC PPR for the same cells shown in ***B***.

### Blocking presynaptic inhibition has no effect on PPR

Paired-pulse responses can also be influenced by presynaptic inhibition. Indeed, under control conditions OSN axon terminals are the site for strong presynaptic inhibition through local glomerular release of both GABA acting on GABA_B_ receptors and dopamine acting on D_2_ receptors (McGann, 2013; Vaaga et al., 2017). To test whether differences in this presynaptic inhibition might contribute to the altered PPR we observed 2 weeks after MMZ treatment, we compared PPR values at both 50 ms and 500 ms ISIs in baseline aCSF versus aCSF containing antagonists for GABA_B_ and D_2_ receptors (Fig 5A,C). In these experiments we again saw an overall effect of increased PPR in the MMZ-treated group for both 50 ms and 500 ms ISIs (Fig 5; 50 ms ISI PPR, PBS: aCSF, 0.67 ± 0.02, n = 43; aCSF + antagonists, 0.65 ± 0.03, n = 15; MMZ(2w): aCSF, 0.76 ± 0.03, n = 27; aCSF + antagonists, 0.70 ± 0.02, n = 20; two-way ANOVA, effect of condition, F_1,101_ = 7.912, P = 0.006; 500 ms ISI PPR, PBS: aCSF, 0.8 ± 0.02, n = 40; aCSF + antagonists, 0.78 ± 0.02, n = 15; MMZ(2w): aCSF, 0.86 ± 0.01, n = 25; aCSF + antagonists, 0.86 ± 0.01, n = 16; two-way ANOVA, effect of condition, F_1,92_ = 14.51, P = 0.0003). However, we observed no effect of the presynaptic receptor antagonists, nor any interaction between MMZ treatment and antagonist application (Fig 5; 50 ms ISI PPR, two-way ANOVA, effect of antagonists, F_1,101_ = 2.824, P = 0.096, effect of interaction, F_1,101_ = 0.80, P = 0.37; Mean diff (predicted), PBS, aCSF vs aCSF + antagonists, +0.02, P = 0.99; MMZ(2w), aCSF vs aCSF + antagonists, +0.07, P = 0.35, Bonferroni’s multiple comparisons test; 500 ms ISI PPR, effect of antagonists, F_1,92_ = 0.254, P = 0.62, effect of interaction, F_1,92_ = 0.441, P = 0.51; Mean diff (predicted), PBS, aCSF vs aCSF+antagonists, +0.02, P > 0.999; MMZ(2w), aCSF vs aCSF+antagonists, +0.003, P > 0.999, Bonferroni’s multiple comparisons test). These data suggest that tonic presynaptic inhibition is minimal in our slice preparation, and that our stimuli were not of sufficient strength to produce phasic presynaptic inhibition over either 50 ms or 500 ms ISIs. More importantly, they show that the higher PPR we observe while regenerating OSNs are re-connecting with OB circuits (Fig 3) is not produced by decreased levels of ongoing presynaptic inhibition.

**Figure 5.**
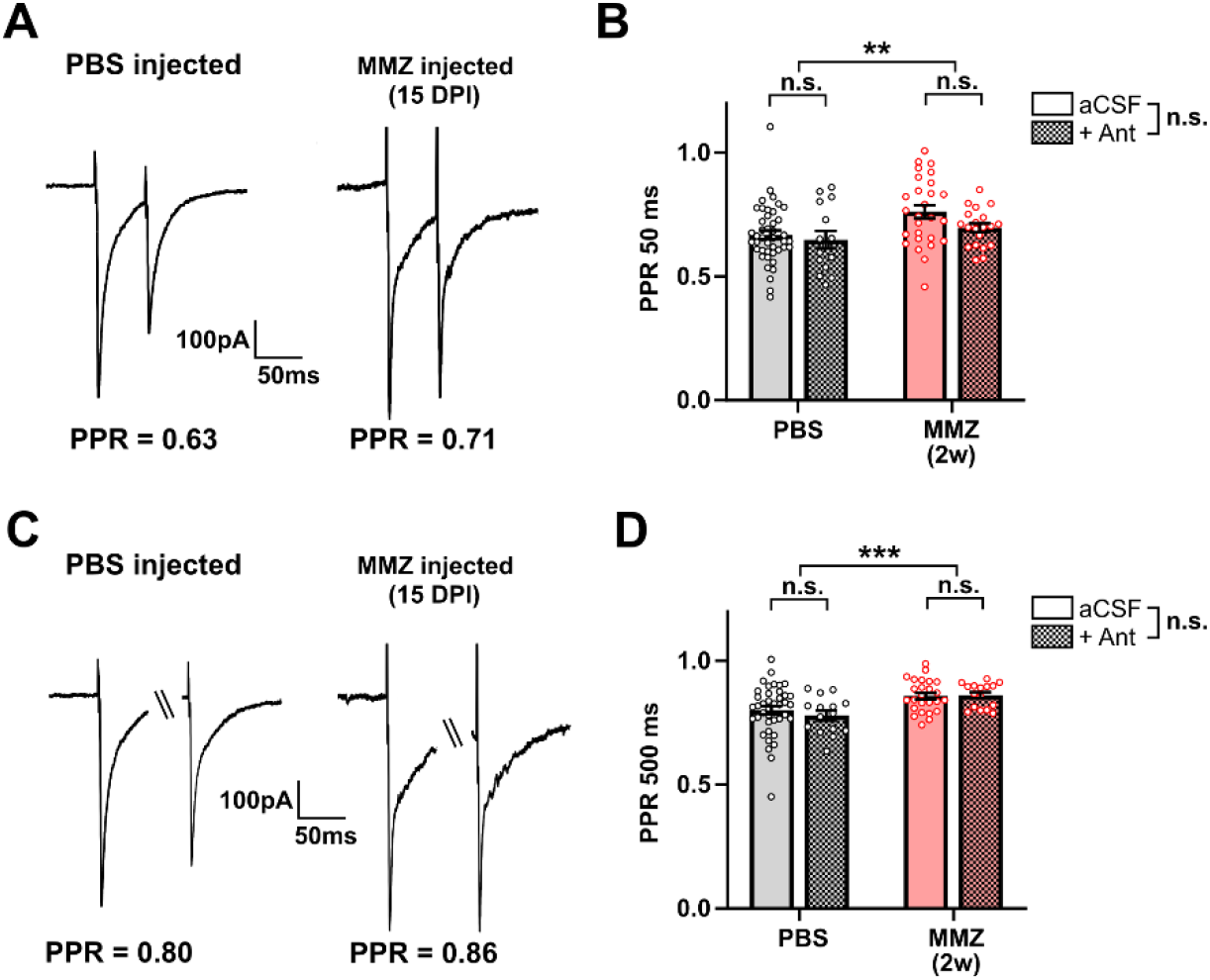
Blocking presynaptic inhibition has no effect on PPR at re-connecting OSN-to-OB synapses. ***A***. Representative postsynaptic currents recorded from an ETC in response to a paired-pulse OSN stimulus with 50 ms inter-stimulus interval (ISI), in a control PBS- (left) or methimazole (MMZ; right)- injected mouse 15 days post injection (DPI). Both recordings were made in the presence of 10μM CGP 55845 and 10μM sulpiride to block GABA_B_ and D_2_ receptors, respectively. PPR, paired-pulse ratio. ***B***. Mean ± SEM PPR at 50 ms ISI recorded in cells (dots) from PBS- or MMZ-injected (2w, 2 weeks post-injection) mice, in control aCSF or in the presence of both GABA_B_ and D_2_ antagonists (+Ant). **, P < 0.01 for the effect of treatment, n.s., non-significant for the overall effect of antagonists or for pairwise tests within treatment groups. ***C***. Representative responses from the same cells shown in A to paired pulse stimuli with 500 ms ISI. Traces have been cropped at the break marks to show individual responses more clearly. ***D***. PPR at 500 ms ISI. ***, P < 0.001 for the effect of treatment; all other conventions as in ***B***.

### An independent measure of decreased release probability in re-connecting terminals

Although PPR is a well-characterised proxy for P_r_ at nose-to-brain synapses (Grubb et al., 2008; Murphy et al., 2004) and was not affected in our data by either post-synaptic desensitisation or presynaptic inhibition, it was nevertheless important to obtain an independent measure of release probability in naturally regenerating OSN-to-OB inputs. We did so by using responses to 50 Hz stimulus trains to calculate the size of the readily releasable pool and therefore directly estimate P_r_ (Fig 6A,B; SMN method; Schneggenburger et al., 1999; Thanawala and Regehr, 2016; Vaaga et al., 2017; see Methods). With this approach we again saw that initially re-connecting OSN inputs at 2 weeks post-MMZ injection had reasonably high P_r_ which was nevertheless significantly lower than in PBS-treated controls (Fig 6C; P_r_, PBS, mean ± SEM, 0.65 ± 0.04, n = 19 cells; MMZ(2w), 0.55 ± 0.03, n = 15 cells; unpaired t test with Welch’s correction, t = 2.2, df = 29.0, P = 0.039). Also in this independent sample, we again found significantly higher 50 ms PPR in MMZ mice (Fig 6D; 50 ms ISI PPR, PBS, mean ± SEM, 0.60 ± 0.02, n = 19 cells; MMZ(2w), 0.71 ± 0.02, n = 15 cells; unpaired t test, t = 3.237, df = 32 P = 0.003), and we observed significant correlations between this PPR measure and P_r_ estimated with the SMN method within both MMZ and PBS treatment groups (Fig 6E; Spearman correlation of P_r_ vs 50ms ISI PPR; PBS, Slope, −0.80 ± 0.36, R^2^ = 0.22, P = 0.041; MMZ(2w), Slope, −0.87 ± 0.24, R^2^ = 0.50, P = 0.003). Two distinct approaches based on paired-pulse and train stimuli therefore both show that during naturally occurring regeneration initially re-connecting OSN inputs have slightly yet significantly lower presynaptic P_r_.

**Figure 6.**
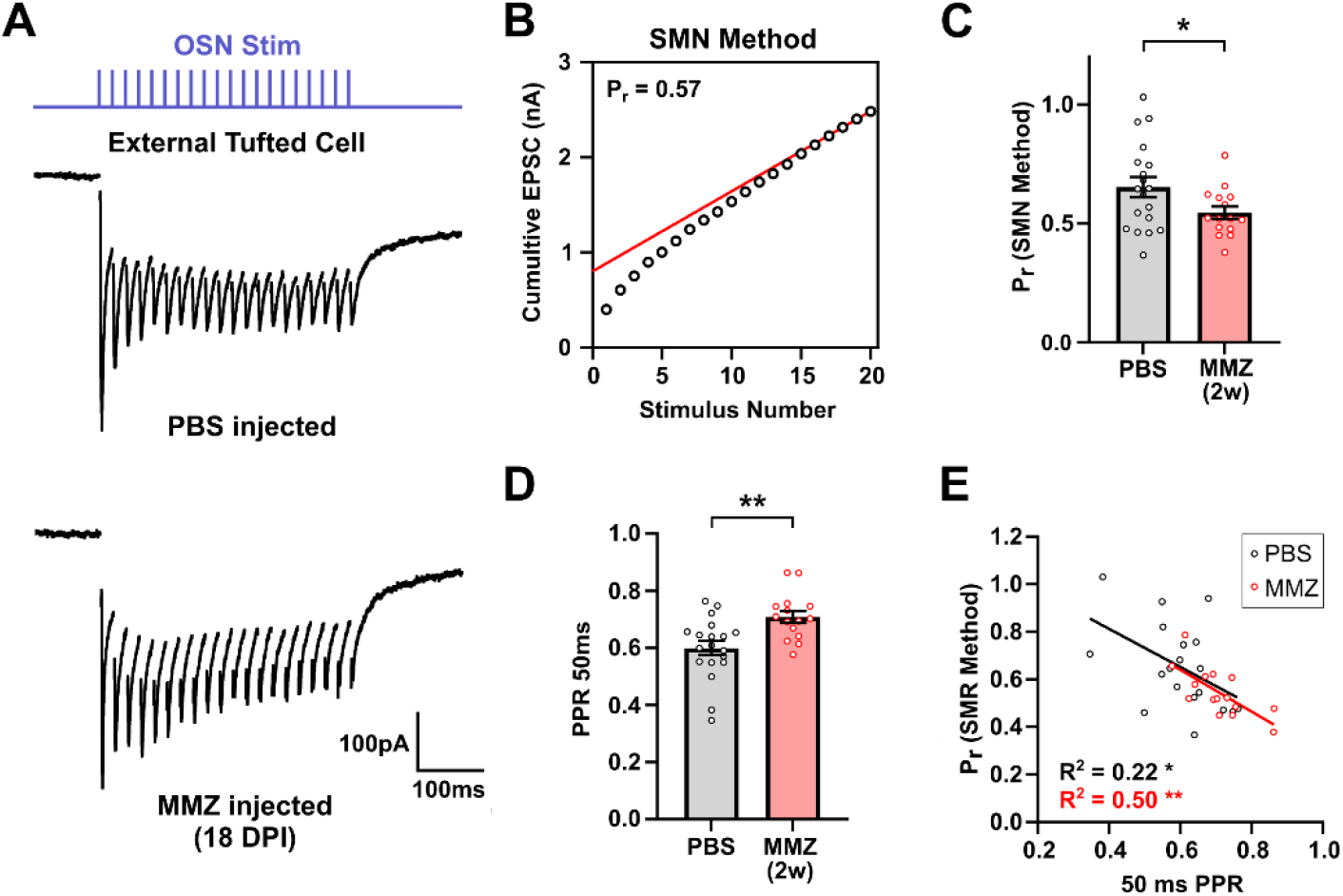
Newly regenerated OSN inputs have higher release probability (P_r_). ***A***. Representative EPSCs recorded in ETCs from PBS- and MMZ-injected mice in response to a 50 Hz train stimulus protocol (Blue trace). ***B***. Calculation of release probability (Pr) based on the Schneggenburger, Meyer, and Neher (SMN) method (see Methods). ***C***. Mean ± SEM of SMN-estimated P_r_ for PBS- and MMZ-treated mice at 2 weeks post-injection (2w). Dots show individual cells; *, P < 0.05. **D.** Mean ± SEM of 50 ms PPR in this independent sample. Dots show individual cells; **, P < 0.01. ***E***. Correlation of SMN-estimated P_r_ with 50 ms ISI PPR. Dots show individual cells; lines show results of best-fit linear regression. *, P < 0.05; **, P < 0.01.

### Newly arriving axon terminals show mature-like sensitivity to D_2_ and GABA_B_ agonists

We found that presynaptic inhibition does not affect regeneration-associated differences in *baseline* PPR (Fig 5), but there is still the possibility that re-connecting OSN terminals might differ from their control counterparts in their *capacity* to undergo GABA_B_- and/or D_2_-based depression. To test whether regenerating OSN axons expressed functional presynaptic receptors, we recorded 50 ms ISI paired-pulse responses in baseline aCSF versus aCSF containing either a D_2_ or GABA_B_ receptor agonist (Fig 7A,D). Both agonists produced a decrease in the amplitude of the first EPSC and an increase in PPR, consistent with a presynaptic effect of reduced P_r_. These effects were indistinguishable in 2 week post-MMZ slices compared to either control or 3 week post-MMZ preparations (Fig 7B,C,E,F; % first peak amplitude after quinpirole application, PBS, mean ± SEM, 46.2 ± 4.9 %, n = 12 cells; MMZ(2w), 34.3 ± 3.6 %, n = 10 cells; MMZ(3w), 45.8 ± 6.6 %, n = 7 cells; one-way ANOVA F = 1.90, P = 0.17; % 50 ms ISI PPR after quinpirole application, PBS, 137.2 ± 7.3 %, n = 12 cells; MMZ(2w), 148.9 ± 9.0 %, n = 10 cells; MMZ(3w), 139.3 ± 9.1 %, n = 7 cells; one-way ANOVA F = 0.59, P = 0.56; % first peak amplitude after baclofen application, PBS, 60.4 ± 5.8 %, n = 15 cells; MMZ(2w), 67.1 ± 10.0 %, n = 7 cells; MMZ(3w), 63.1 ± 9.3 %, n = 8 cells; one-way ANOVA F = 0.18, P = 0.84; % 50 ms ISI PPR after baclofen application, PBS, 128.1 ± 6.5 %, n = 15 cells; MMZ(2w), 128.8 ± 10.8 %, n = 7 cells; MMZ(3w), 140.4 ± 17.2 %, n = 8 cells; Kruskal-Wallis test, K-W statistic = 0.372, P = 0.83). Even at the very first stages of re-connection with downstream circuits, then, regenerating OSN axon terminals have a fully mature-like capacity for presynaptic inhibition via both GABA_B_ and D_2_ receptor activation.

**Figure 7.**
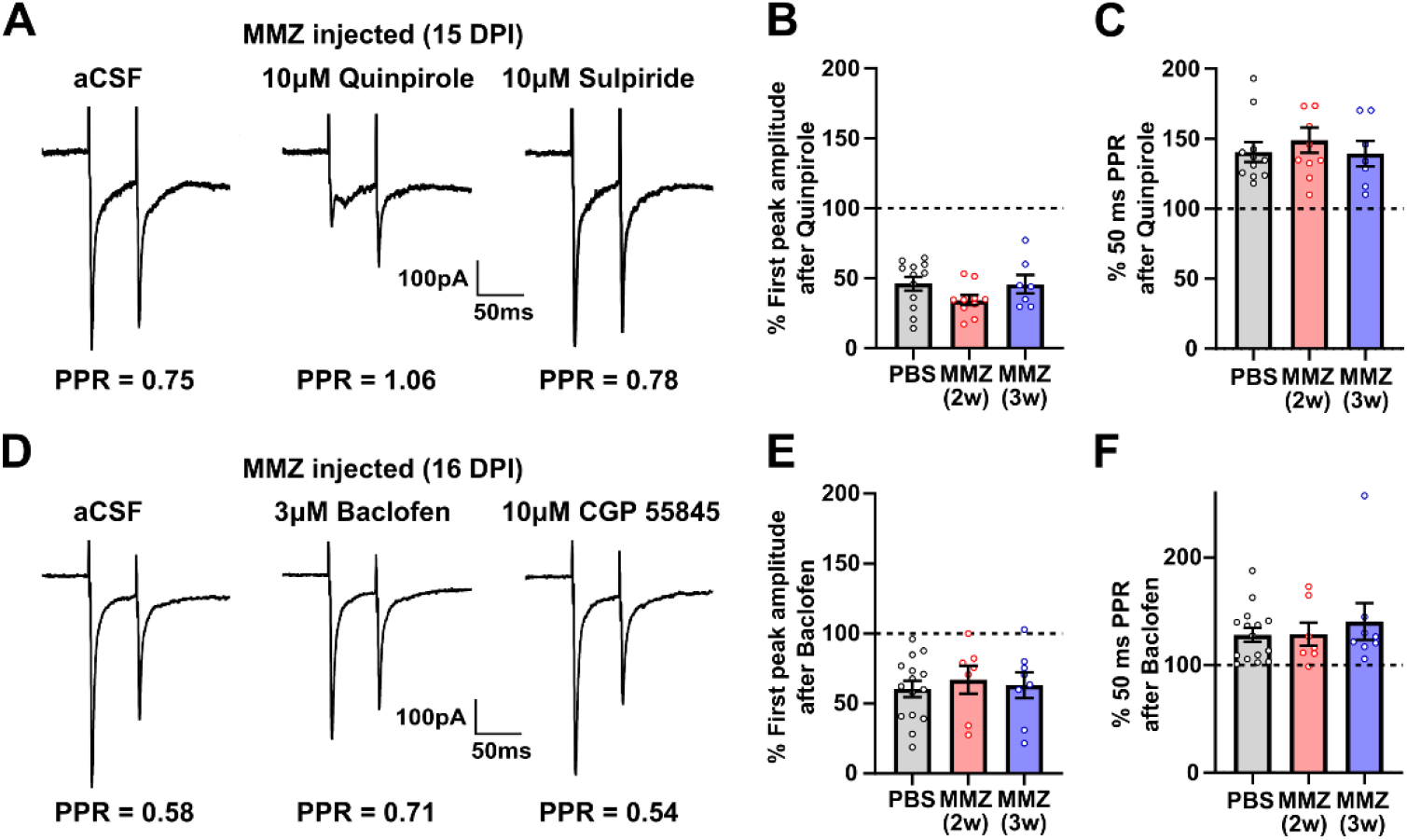
Newly arriving axon terminals show mature-like sensitivity to D_2_ and GABA_B_ agonists. ***A***. EPSCs recorded in an ETC from a methimazole (MMZ)-treated mouse (15 days post-injection) in response to a paired-pulse OSN stimulus with 50 ms inter-stimulus interval (ISI), in the absence (aCSF; Left), and presence of sequentially added quinpirole (10μM, Middle) and sulpiride (10μM, Right). Application of the D_2_ agonist quinpirole resulted in a decrease in the first evoked postsynaptic current amplitude and a concomitant rise in paired-pulse ratio (PPR). Both effects were reversed by the D_2_ antagonist sulpiride. ***B***. Mean ± SEM change in first peak amplitude after application of quinpirole. ***C***. Mean ± SEM change in PPR after application of quinpirole. ***D***. ETC responses to 50 ms ISI paired-pulse OSN stimulation in the absence (aCSF; Left), and presence of sequentially added baclofen (3μM, Middle) and CGP 55845 (10μM, Right). Application of the GABA_B_ agonist baclofen resulted in a decrease in the first evoked postsynaptic current amplitude and a rise in PPR. Both effects were blocked by the GABA_B_ antagonist CGP 55845. ***E***. Mean ± SEM change in first peak amplitude after application of baclofen. ***F***. Mean ± SEM change in PPR after application of baclofen. In ***B, C, E & F***, dots show data from individual cells; 2w, 2 weeks post-MMZ; 3w, 3 weeks post-MMZ.

## Discussion

We have found that naturally regenerating OSN axons begin to re-establish synaptic input onto central targets around 2 weeks after olfactotoxin-induced degeneration. Starting from this 2-week time point, functional connectivity with post-synaptic OB neurons is rapidly developed, and presynaptic properties at these re-connecting inputs are almost fully mature. However, rapid maturation is seen in the paired-pulse ratios of OSN-evoked monosynaptic EPSCs, which are significantly and briefly higher in the third week post-MMZ treatment, before reaching control levels around one week later. This maturational effect could not be explained by differences in postsynaptic receptor desensitisation nor tonic presynaptic inhibition, and – because it was mirrored by an independently measured decrease in release probability – likely reflects transiently immature intrinsic presynaptic properties. At the same time, we found that even the most immature re-connecting terminals have a fully mature capacity for presynaptic inhibition via both GABAergic GABA_B_ and dopaminergic D_2_ receptor activation.

### Presynaptic maturation at developing and regenerating synapses

Across the nervous system, synaptic development is characterised by large variability in presynaptic maturational processes. At many connections forming for the first time, PPR and/or P_r_ show no developmental changes at all (Bornschein et al., 2019; Ferrer et al., 2018; Fukuda et al., 1993; Grubb et al., 2008; Hsia et al., 1998; Speed and Dobrunz, 2009). At others, including most excitatory cortical synapses and the highly specialised Calyx of Held, presynaptic maturation is accompanied by developmental increases in PPR and/or decreases in P_r_ (Bolshakov and Siegelbaum, 1995; Choi and Lovinger, 1997; Feldmeyer and Radnikow, 2009; Pouzat and Hestrin, 1997; Reyes and Sakmann, 1999; Speed and Dobrunz, 2009; Taschenberger and von Gersdorff, 2000), although in some cases these effects can be offset by other presynaptic changes to produce an overall increase in synaptic fidelity with age (Taschenberger and von Gersdorff, 2000). Our present results are most consistent with effects found at other developing synapses, including parallel fibre inputs onto cerebellar Purkinje cells and GABAergic inputs from the hippocampal dentate gyrus to CA3 pyramidal cells, where PPR can decrease, and/or P_r_ can increase across maturation (Baur et al., 2015; Caillard et al., 1998; Dietz et al., 2011; Katayama et al., 2017; Mori-Kawakami et al., 2003). Our data showing fully functional presynaptic receptors at immature axon terminals are also consistent with similar observations at various developing synapses (Bosch and Ehrlich, 2015; Caillard et al., 1998; Fukuda et al., 1993; Mizutani et al., 2006).

At regenerating synapses, by far the best-characterised process of functional presynaptic maturation is at the vertebrate neuromuscular junction. There, re-connectivity is associated with an overall improvement in presynaptic function over time, with increasing action potential invasion (Dennis and Miledi, 1974; Miledi, 1960), increasing quantal content (Argentieri et al., 1992; Decino, 1981; Dennis and Miledi, 1974), increasing P_r_ (Argentieri et al., 1992) and increasing structural organisation at the active zone (Ko, 1984) all contributing to the return to pre-injury levels of synaptic fidelity. Such effects are consistent with the rapid maturation towards control levels of PPR that we observed at regenerating olfactory nerve synapses (Fig 3), as is the finding that fully mature regenerated inputs in the adult lamprey spinal cord have P_r_ values equal to those in uninjured animals (Parker, 2017). In terms of presynaptic inhibition, our data (Fig 7) are consistent with one other observation of presynaptic receptors being present and functional at regenerating central axon terminals – in the re-connecting post-injury goldfish retinotectal projection, the presynaptic inhibitory effects of adenosine are just as strong as in control preparations (Zhang and Schmidt, 1998). However, more work on the maturation of different natural and experimentally induced models of synaptic regeneration is required to see whether there are exceptions to these seemingly consistent findings.

### Presynaptic maturation at the first synapse in olfaction

Our finding of functional presynaptic receptors at initially re-connecting OSN-to-OB synapses (Fig 7) is also consistent with evidence for mature levels of GABAB-mediated presynaptic inhibition at this connection in early postnatal development (Grubb et al., 2008). In contrast to our present results (Fig 3), however, neonatal nose-to-brain synapses showed completely mature-like PPR from the very earliest ages studied, with no evidence for any maturational changes across the first postnatal month (Grubb et al., 2008). This could potentially suggest that mechanisms of presynaptic maturation are different during early development versus regeneration of the olfactory nerve. However, the earliest functional recordings of OSN-to-OB inputs took place at postnatal day(P) 1 (Grubb et al., 2008), while ultrastructural evidence suggests that the first synaptic connections can occur between OSN axon terminals and OB as early as embryonic day (E) 14-15 (Blanchart et al., 2008; Hinds and Hinds, 1976; Tufo et al., 2022). Moreover, the molecular and anatomical maturation of OSNs occurs at a slower pace in later postnatal life versus early development (Liberia et al., 2019). It is possible, then, that development and regeneration share functional features, and that a short, transient period of higher PPR values could also occur during the initial pre-natal stages of OSN-to-OB synaptic formation.

The functional properties and potential maturation of immature presynaptic terminals have not yet been assessed during baseline conditions of *constitutive,* ongoing OSN neurogenesis. However, the synaptic boutons of immature OSNs in the adult OB are known to be highly structurally dynamic, to make functional monosynaptic connections with OB neurons, and – crucially – to be ultrastructurally indistinguishable from those of neighbouring, mature OSNs (Cheetham et al., 2016; Huang et al., 2021). This could potentially indicate a lack of presynaptic maturation in constitutively generated OSNs, and therefore a fundamental difference between OSN terminals connecting under constitutive or regenerative conditions. Alternatively the ultrastructural parity observed between constitutively generated immature and mature OSNs (Cheetham et al., 2016) could hide significant functional distinctions, especially if – as we observe here – those functional differences are between quantitatively different amounts of overall high release probability. If so, nose-to-brain synaptic maturation might follow a similar trajectory in both constitutive and regenerative modes of OSN neurogenesis. Electrophysiological recordings of PPR evoked from immature OSNs in uninjured mice are needed to address this question.

Such experiments would also aid interpretation of another distinction. Unlike the rapid maturation of PPR seen here in regenerating OSN inputs (Fig 3), no maturational changes in PPR were observed in OSN inputs onto adult-born OB neurons. Instead, newly connected adult-generated OB cells could receive OSN inputs with fully mature levels of strong paired-pulse depression (Grubb et al., 2008). If inputs from constitutively generated immature OSNs *do* have transiently higher PPR values, like those of initially re-connecting regenerating OSNs (Fig 3), this would indicate that adult-born OB neurons *do not* preferentially receive input from immature OSNs at any stage in their maturation. At the one point in the mammalian brain where immature adult-born pre- and postsynaptic neurons might come into contact, then, they would not necessarily have a preference for connecting with each other.

Finally, our data showing maturational changes in presynaptic function at nose-to-brain synapses are fully consistent with previous studies demonstrating considerable presynaptic plasticity at this first synapse in olfaction. Release probability at OSN terminals is increased by sensory deprivation (Tyler et al., 2007). Such homeostatic plasticity could potentially also be at play in re-connecting axons, if overall activity levels in newly generated OSN were to decrease – and thus trigger a compensatory P_r_ increase – as they mature. OSN presynaptic function can also be dynamically regulated by learning (Bhattarai et al., 2020; Kass et al., 2013) and by longer-term alterations in the olfactory environment (Tsukahara et al., 2021). Given that sensory experience can regulate the anatomical regrowth of OSN axons after MMZ-induced degeneration (Kikuta et al., 2015), it would be fascinating to know whether such plastic processes might also regulate the rapid functional maturation we observed in regenerating OSN-to-OB inputs.

### Implications of presynaptic maturation for functional axon regeneration

At mature OSN terminals, functional specialisations including high P_r_ have been proposed to contribute to the reliable detection of even weak odorant stimuli (Murphy et al., 2004). In regenerating conditions, where an animal’s survival might depend on their ability to send odorant-evoked information across a few initially formed synaptic connections, it is therefore unsurprising that presynaptic function is almost fully mature upon first contact, and that it develops so rapidly thereafter. However, it might still seem maladaptive to have even a brief phase of immature presynaptic function as connectivity is first re-established. Why are regenerating OSN terminals not completely mature and ready to ‘plug and play’ as soon as they reach the OB? One potential reason could be the apparent incompatibility seen elsewhere in the regenerating nervous system between axon growth and presynaptic maturation. After spinal cord injury in mice, several molecular components of presynaptic release machinery can act as a brake on re-growth in primary sensory axons (Hilton et al., 2021), suggesting that in order to get re-growing axons to their targets, full presynaptic function must be suppressed until the point of synaptic contact. Indeed, genes encoding presynaptic proteins are some of the last to be expressed in OSN development, at least under constitutive conditions (Marcucci et al., 2009; McClintock et al., 2020). Alternatively, the briefly higher PPR at immature re-connecting OSN inputs could actually be adaptive in its own right. Depending on the patterns of odorant-evoked spiking present in immature OSNs – which are currently entirely unknown – less depression of closely-spaced EPSCs could contribute to higher fidelity of information transfer across the first synapse in this sensory system. Finally, in a manner non-mutually exclusive from either of those possibilities, any transient immaturity in presynaptic function could be offset by plastic changes in downstream circuits in the OB or higher olfactory processing centres. If postsynaptic mechanisms can compensate for brief periods of intrinsic presynaptic immaturity and/or the presence of fully functional receptors for presynaptic inhibition, even immature regenerating inputs to the brain might still be able to drive adaptive sensory-evoked behaviour (Huang et al., 2021).

## Methods

### Animals

All experiments were carried out using adult (≥ 2 months old) wild-type C57/Bl6J mice (Charles River) of either sex, housed under a 12 h light-dark cycle with access to water and food *ad libitum.* All animal work conformed to United Kingdom legislation outlined in the Animals (Scientific Procedures) Act 1986. All experiments were performed under UK Home Office personal and project licences held by the authors.

### Methimazole administration

Methimazole (MMZ; Sigma M8506) was made up to 10 mg/ml in PBS. Mice received a single intraperitoneal injection of either PBS (control) or 100 mg/kg MMZ. Mice treated with MMZ typically experienced mild, temporary loss of ~10 % of their initial weight and/or reduced mobility over the first 1-2 d post-injection, so were given wet food and/or time in a heated recovery chamber on a discretionary basis.

### Immunohistochemistry

Mice were anaesthetised with an intraperitoneal overdose of pentobarbital and then perfused with 20 ml PBS with heparin (20 units/ml; Alfa Aesar A16198), followed by 20 ml of 4 % paraformaldehyde (PFA, TAAB Laboratories, P001; in 60 mM PIPES, 25 mM HEPES, 5 mM EGTA, and 1 mM MgCl_2_).

To expose the olfactory epithelia, the rostral half of the calvaria (anterior to bregma) was removed, and the samples were first postfixed overnight in 4 % PFA (4°C) and then placed in 0.25 M EDTA (Invitrogen AM9261) in PBS at 4°C for 3 d for decalcification. Samples were washed in increasing concentrations of sucrose in PBS (10 %, 20 % and 30 %; Sigma Millipore, S9378), and kept overnight at 4°C in 30 % sucrose. Samples were then embedded in OCT (VWR Chemicals, 00411243), frozen in liquid nitrogen, and sectioned coronally on a cryostat (Leica Microsystems, CM 1950) into 50 μm slices. Olfactory bulbs were dissected and postfixed in 4 % PFA for 24 h, then the right OB from each mouse was embedded in 6 % agarose and sliced coronally at 50 μm using a vibratome (Leica VT1000S).

For OMP fluorescent immunohistochemistry, slide-mounted OE sections or free-floating OB slices were washed with PBS and incubated in 3 % BSA (Merck, A2153) with 0.25 % triton and 0.02 % sodium azide (Severn Biotech Ltd, 40-2000-01) for 2 h at room temperature. They were then incubated in primary antibody solution, consisting of 1:10,000 goat anti-OMP antibody (Wako, 544-10001) in PBS with 3 % BSA, 0.25 % triton and 0.02 % sodium azide for 2 d at 4°C. After washing three times for 5 min with PBS, sections were incubated in secondary antibody solution, consisting of 1:1000 Alexa Fluor 488 donkey anti-goat IgG (ThermoFisher A11055) in PBS with 3 % BSA, 0.25 % triton and 0.02 % sodium azide, for 2 h at room temperature. After washing again three times for 5 min with PBS, samples were incubated with NucRed Live 647 (Invitrogen, R37106), in PBS for 30 min at room temperature. After one last wash of 5 min in PBS, free-floating slices were then mounted on glass slides, and all sections were coverslipped with FluorSave Reagent (Millipore, 345789).

For VGlut2 labelling of free-floating OB sections (with 1:10,000 guinea pig anti-VGlut2, Sigma AB2251-I; and 1:1000 Alexa Fluor 568 goat anti-guinea pig IgG, ThermoFisher A11075), we followed the above procedure exactly, with the exception that all blocking and antibody incubation steps took place in 5 % normal goat serum (NGS) in PBS with 0.25 % triton and 0.02 % sodium azide.

### Image acquisition and analysis

All image acquisition and analysis was carried out blind to experimental group. Images were acquired with a laser scanning confocal microscope (Zeiss, LSM 710) using appropriate excitation and emission filters, a pinhole of 1 AU and a 40x oil immersion objective. Laser power and gain were set to avoid signal saturation, and were kept consistent across slices from PBS- and MMZ-injected mice fixed at the same timepoint. For OMP label in the olfactory epithelium, a Z-stack (dimensions: 512×512 Pixels, 212.55 x 212.55 μM, pixel size 0.415 μM^2^, z-step: 1 μM) covering the full depth of the tissue was acquired from each of four different epithelial regions on each section (two septal, and two in the turbinates). For OMP and VGlut2 label in the OB, NucRed nuclear label was used to guide acquisition of single-plane images of randomly chosen glomeruli (dimensions: 512×512 Pixels, 141.70 x 141.70 μM pixel size 0.277 μM^2^), at the z-plane where their area was maximal.

We used ImageJ to calculate OSN density in each z-stack image from the olfactory epithelium, from the z-plane with the highest OMP+ cell number. We drew a line parallel to the lamina propria through the densest region of OMP staining, counted all OMP+ cells in contact with this line, and divided by line length to obtain an absolute OSN density value in cells / μm. To standardise across samples that had been fixed at different times, we took advantage of the fact that all MMZ-treated mice that were fixed on a given day were always fixed and processed alongside a PBS-injected control. We therefore normalised MMZ-treated OSN density values by dividing by the mean OSN density value from co-fixed PBS-treated tissue (1 = control level).

Glomerular label in the OB was analysed with custom routines in Matlab, using a simple threshold-based algorithm which standardised for potential staining differences across slices by normalising fluorescence intensity *within* each image (Kikuta et al., 2015). Background fluorescence was determined for each image by taking the mean grey value of a small, manually selected region of the OB’s external plexiform layer. The threshold for ‘positive’ pixels was then set at three times this background value. After drawing an ROI of the glomerular neuropil in the NucRed channel, the percentage was then calculated of ‘positive’ (i.e. ≥ 3 x background) pixels within this area.

### Acute slice electrophysiology

Mice were deeply anaesthetised under isoflurane before being decapitated. The OBs were rapidly dissected and transferred into ice-cold slicing medium containing (in mM): 240 sucrose, 5 KCl, 1.25 Na_2_HPO_4_, 2 MgSO_4_, 1 CaCl_2_, 26 NaHCO_3_ and 10 D-Glucose, bubbled with 95 % O_2_ and 5 % CO_2_. 300 μm horizontal sections were cut using a vibratome (Leica, VT1000S) and slices were maintained at 32°C for 30 min in artificial cerebrospinal fluid (aCSF) containing (in mM): 124 NaCl, 5 KCl, 1.25 Na_2_HPO_4_, 2 MgSO_4_, 2 CaCl_2_, 26 NaHCO_3_ and 20 D-Glucose, bubbled with 95 % O_2_ and 5 % CO_2_; pH 7.3-7.4, Osmolarity: 300-310 MOsm). Slices were then transferred to room temperature aCSF for 30 min before experiments began.

For whole-cell patch-clamp recordings, individual slices were placed in a chamber mounted on a Nikon FN1 Fixed Stage Microscope and held in place with a stainless-steel anchor. OB slices were perfused with aCSF heated to 32–34°C with an in-line heater (TC-344B, Warner), and were visualised using a 40x water immersion objective. ETCs were identified by their location (near the border between the glomerular layer and external plexiform layer), shape (balloon-shaped soma around 10 μm in diameter) and distinctive apical dendrite ramifying within a single glomerulus (Hayar, et al., 2004; Galliano et al., 2021). Glass recording electrodes (outer diameter 1.5 mm, inner diameter 0.84 mm, World Precision Instruments 1B150F-4) were pulled using a vertical puller (Narishige, PC-10) and had a resistance of 3-6 MΩ when filled with an intracellular solution containing (in mM): 130 CsMeSO_4_, 4 MgCl_2_, 0.2 EGTA, 10 HEPES, 10 Na-Phosphocreatine, 4 Na_2_ATP, 0.4 Na_3_GTP, alexa 488 (pHed to 7.2 with CsOH; ~290 MOsm; uncorrected liquid junction potential ~10.4 mV). Fluorescent label of Alexa-filled neuronal morphology was visualised using LED excitation (CoolLED pE-100) with appropriate excitation and emission filters. Recordings were performed using a Heka EPC10/2 amplifier with PatchMaster acquisition software. Signals were digitised and sampled at 20 kHz (50 μs interval sample), and Bessel filtered at 10 kHz (filter 1) and 2.9 kHz (filter 2). Data were excluded if series resistance (Rs), measured from the peak current to a 10 mV hyperpolarising voltage step in voltage-clamp, exceeded 30 MΩ or varied by > 20 % over the course of the recording.

OSN evoked responses were produced by electrically stimulating OSN fibres using a monopolar patch-pipette filled with aCSF solution (see above) placed into the olfactory nerve layer upstream of the target glomerulus. 1 ms current pulses were applied by an isolated stimulator (0.01-10mA, ISO-Flex, A.M.P.I.), with amplitudes ranging from 0 to 80 μA. At amplitudes of > 100 μA it was possible to produce direct stimulation of the glomerular components, as evidenced by observations of post-synaptic currents even in the presence of blocking concentrations of TTX. Monosynaptic EPSCs were identified by their fast, reliably timed onset latency (< 10 ms), and slower offset kinetics (see Figure 2Ci), and stimulation intensity was set to produce reliable EPSC responses of amplitude 200-900 pA. All drugs used for electrophysiological experiments were acquired from Sigma-Aldrich (Merck), dissolved in aCSF from frozen stocks, and added to the slice via perfusion. Complete dialysis of the bath chamber with drug solution occurred in ~ 2.5 min post-addition with post-drug recordings made ≥5 min post-addition.

### Analysis of electrophysiological parameters

Exported traces were analysed using custom written routines in Matlab (Mathworks). ETC membrane properties were estimated from a 10 ms, 10 mV hyperpolarizing voltage step from −60mV in voltage clamp. The resulting current transients were analysed to provide measures of series resistance (Δvoltage step/Δpeak current), input resistance (Δvoltage step/Δsteady-state current) and cell capacitance (integral of the area under the current transient). All reported passive membrane properties were measured immediately after membrane rupture.

Paired-pulse responses were measured from postsynaptic ETCs by stimulating tracts of nearby OSNs twice in succession with an inter-stimulus interval of either 50 ms or 500 ms. The relative ratio of the two peaks’ amplitude was then calculated to give the paired-pulse ratio (PPR). The first response amplitude was measured from baseline current before stimulus onset to the maximum current peak of the first stimulus response. Because of the relative slow EPSC offset kinetics, the baseline for the second response was obtained by using a fitted single exponential function to extrapolate the decay of the first response until the peak of the second response. This was then used as the baseline from which to calculate the second response amplitude.

To estimate P_r_ with the SMN method (Schneggenburger et al., 1999; Thanawala and Regehr, 2016; Vaaga et al., 2017), OSNs were stimulated at a frequency of 50Hz with 20x 1 ms pulses whilst post-synaptic recordings were made from downstream ETCs. EPSC amplitudes were measured using baselines obtained from exponential fits to each preceding event, as for PPR measures (see above). A linear fit was applied to the final 5 EPSCs on a plot displaying cumulative EPSC amplitude versus stimulus number. The y-intercept of the fitted line was taken as an estimate of the readily releasable pool (Schneggenburger et al., 2002). P_r_ was then calculated by dividing the amplitude of the first evoked EPSC by the size of the readily releasable pool. An interval of at least one minute was given before the beginning of each stimulation protocol to ensure the complete replenishment of vesicle pools. In each case at least 2 repetitions, but typically ≥ 3 repetitions of each protocol were run, and the reported single cell values represent an average measure of these P_r_ estimates in each cell.

### Statistical analysis

Statistical analysis was carried out using Prism (GraphPad, San Diego, USA), and details of all individual statistical tests are reported alongside the appropriate results. α values were set to 0.05 and all comparisons were two tailed. Distributions were assessed for normality using the D’Agostino and Pearson omnibus test, and parametric or non-parametric tests performed accordingly.

## Author Contributions

LPB, AC and MSG designed experiments; LPB and AC performed experiments; LPB, AC and MSG analysed data; LPB and MG wrote the paper.

## Acknowledgements

This work was supported by grants to MSG from the International Foundation for Research in Paraplegia (P160), European Research Council (FUNCOPLAN; 725729), and Biotechnology and Biological Sciences Research Council (BB/V000195/1). We thank Christiane Hahn for protocol optimisation, and Juan Burrone for comments on the manuscript.

